# LINE-1 ORF1p is a shared and immunogenic antigen in cancer

**DOI:** 10.1101/2025.06.11.659092

**Authors:** Wilson McKerrow, Megan Snyder, Chong Chu, Deena Kelly, Heike Keilhack, Liyang Diao

**Affiliations:** ROME Therapeutics, Inc

**Keywords:** Cancer vaccine, immunopeptidomics, LINE-1, shared cancer antigen

## Abstract

**Background:** Recent clinical trials are beginning to show the potential for therapeutic cancer vaccines directed against tumor associated antigens. However, there is a limited set of well-defined antigens that are shared across tumors. Noncanonical coding elements in the genome that are usually silenced in normal cells potentially represent a rich source of “dark” antigens when expressed and presented to the immune system. LINE-1 elements are repetitive, virus-like genomic sequences that can copy themselves to new genomic loci, and which are reactivated in many cancers. In this study, we explore ORF1p, a protein encoded by LINE-1 elements, as a potential shared cancer vaccine antigen.

**Methods:** We assessed the tumor specificity of LINE-1 expression in large scale public RNA-seq data sets and in a curated database of public immunopeptidomics studies. We validated these findings using cell line and tissue data and performed an *in vitro* vaccination assay using healthy donor PBMCs to test the immunogenicity of ORF1p.

**Results:** We found widespread RNA expression of LINE-1 across many tumor types with esophageal cancer showing the most highly elevated expression. We also found widespread MHC class I presentation of ORF1p-derived peptides in a range of tumor types, including esophageal cancer. We validated the tumor specific protein expression of ORF1p in a small set of 5 tumor, 5 matched normal, and 5 healthy esophageal samples. Finally, we demonstrated that dendritic cells pulsed with ORF1p peptides were able to induce interferon gamma production in healthy donor human T cells after repeated stimulation *in vitro*.

**Conclusions:** We present data demonstrating that dark antigens derived from LINE-1 elements are 1) expressed in a cancer tissue-specific manner, 2) detected in immunopeptidomics data (both publicly available and newly generated for this study) and 3) immunogenic *in vitro*. The tumor specificity and immunogenicity of ORF1p peptides point to their potential as a cancer vaccine antigen.

## Background

Numerous recent clinical trials have shown the therapeutic potential for cancer vaccines (1–6). While there are a range of delivery platforms (1,3–5,7,8), they all seek to elicit a T cell immune response against an antigen that is specifically, or at least differentially, expressed in cancer cells. Most recent clinical successes have focused on “neoantigens” – mutations that occur within the tumor that give rise to non-self epitopes (9). While there are some recurrent mutations, such as certain KRAS mutations (10), most tumor mutations are specific to the tumor in which they arose and thus require a “personalized” cancer vaccine that is specific to that patient’s tumor. Another strategy is to target so-called tumor-associated antigens (TAAs). These molecules, such as those encoded by PRAME and the MAGEA family, are preferentially expressed in tumors, and, despite being encoded in the genomes of every cell, are capable of eliciting a T cell response (1). However, the number of such TAA-encoding genes is limited. For a cancer vaccine to reach a large number of patients, it will require novel classes of immunogenic TAAs that are highly shared across tumors.

Transposable elements, which constitute half of the human genome and can copy themselves to new locations, are a potential source of novel antigens (11). Because many of them are fixed in the human population, they can be highly shared. At the same time, they have no known function that benefits their human host and instead act as genomic parasites. Residing within this gray area between self and non-self may make these elements more immunogenic than traditional TAAs. Moreover, given these repetitive elements are expressed from multiple genomic loci, they would be resistant to immune evasion through antigen down-regulation, as that would require the tumor to simultaneously modulate dozens or more loci. This offers an important advantage compared to TAAs encoded by conventional genes.

In humans, Long Interspersed Nuclear Element-1 (LINE-1) is the only protein-coding transposable element that remains transpositionally active (12). We recently showed that LINE-1 elements are highly expressed in a range of tumor types and can make dozens or more somatic insertions in a single tumor (13). LINE-1 encodes two proteins, ORF1p and ORF2p, that are involved in making new copies of itself. Of these, ORF1p is more much abundantly expressed (14), and has been shown to have a tumor specific expression pattern in several studies (15,16). Antibodies targeting ORF1p have been found in cancer patients (17) and in Lupus patients (18), but little is known about how T cells, which are key to vaccine efficacy, might respond to this antigen.

We tested the potential for ORF1p as a cancer vaccine target, first validating its tumor specific expression at the RNA level. To determine whether ORF1p is presented on major histocompatibility complex (MHC) class I molecules, and therefore visible to the immune system, we analyzed public immunopeptidomics data from 47 studies and identified epitopes that span an array of HLA alleles and tumor types. We also showed that ORF1p elicits a T cell response *in vitro*.

## Methods

See Extended Methods for more detail.

### Quantification of LINE-1 from RNA-seq

LINE-1 quantification analyses were performed as described previously (13). Briefly, RNA-seq reads from The Cancer Genome Atlas (TCGA) (19), the Genotype-Tissue Expression (GTEx) (19), and the Cancer Cell Line Encyclopedia (CCLE) (20) were aligned using the STAR aligner (21). Locus specific LINE-1 expression was quantified using L1EM (22) and converted to transcripts per million (TPM), which were summed across full length intact loci as annotated in L1Base (23). These are N=146 loci curated to be full length and have both intact ORF1 and ORF2 proteins in humans.

### Cancer Tissue Preparation

Frozen tissues from 5 esophageal tumors, 5 non-malignant adjacent, and 5 healthy esophageal samples were obtained from BioIVT and pulverized using the Covaris cryoPREP instrument. Full sample details can be found in Supplemental Table 1.

### Protein Simple Western blot (Jess instrument)

ORF1p Western blots (excluding CCLE cell lines shown in Figures 2 C, D) were performed using the Protein Simple Jess Automated Western Blot System, using a commercially available rabbit monoclonal antibody (ab246317 from Abcam). Bands from 39kd to 50kd were considered ORF1p. ORF1p has an apparent molecular weight of 40 kd with traditional Western blotting (24) but can appear at higher molecular weights using the Protein Simple Western blotting technique, depending on the cell or tissue extraction buffer and methods. The corrected area (corr. area) was used for quantification and values were normalized to 2 µg of total loaded protein.

### Traditional Western (CCLE cell lines)

ORF1p quantification in CCLE cells was performed at Pharmaron using a standard Western blot with a commercially available ORF1 antibody (Sigma: MABC1152). Quantifications were normalized to beta-actin.

### HeLa overexpression systems

To generate stable HeLa cell lines expressing a codon optimized human LINE-1 element, DOX-inducible L1 ORFeus and C-terminally tagged HLA-A*0201 piggyBac-compatible plasmids were transfected into cells along with the piggyBac transposase. The cells were then selected using Hygromycin (100 µg/ml) and Puromycin (0.5 µg/ml) for one week.

### Immunopeptidomics experiments

Immunopeptidomics was performed by Biognosys using the TrueDiscovery® platform. Samples were lysed and MHC class I complexes were immunoprecipitated using the W6/32 antibody and peptides were eluted from the complexes. Tissue samples were split into 5 fractions; cell line samples were not fractionated. Peptides were then loaded on a liquid chromatography column and analyzed by data dependent acquisition (DDA) mass spectrometry. Peptide spectral matches (PSMs) at a false discovery rate (FDR) of <1% were identified by the Spectronaut software. For the esophageal tumor samples, the database search was supplemented by ORF1p peptide sequences translated from loci that have an uninterrupted ORF1 and are expressed at least 2 TPM (L1EM estimate) in at least one of the tumor samples.

### Analysis of public immunopeptidomics data

Raw files were downloaded from the relevant repository and then converted to mzml using msconvert. Comet (25) was used to query mass spectra against the uniprot human proteome. Decoy search was set to concatenated search, peptide mass tolerance was set to 20 ppm, fragment bin tolerance was set to 0.02 Daltons (Da), search enzyme was set to none, allowed peptide length was 8-11, oxidation (15.9949 Da) of methionine was a variable modification, and carboxyamidomethylation (57.021464 Da) of cysteine was a fixed modification. The comet matches were then rescored using the Prosit (26) model as implemented in Oktoberfest (27). ORF1p hits across all runs were merged into a single table and matches at an FDR of 1% were identified using a semi-supervised SVM strategy as outlined by percolator (28). Binding predictions were assessed using MHCflurry with default parameters (29).

### Generation of ORF1p peptides for T cell immunogenicity assay

Using the canonical peptide sequence of ORF1p (L1RE1 in uniprot), we generated a list of 82 15-mer peptides with 11 bp overlaps. Peptides were synthesized by GenScript Biotech (Piscataway, NJ). Two peptides with low purity (<90%) were excluded. Peptides were pooled into eight groups of ten peptides, in sequential order, starting at the N’ terminus of ORF1p. A positive control consisting of 27 peptides from common pathogens (Clostridium tetani, Epstein-Barr Virus (EBV), Human Cytomegalovirus (HCMV), Influenza A; also known as the CEFT pool) and a negative control consisting of peptides from the human MOG gene (Myelin Oligodendrocyte Glycoprotein; also known as the MOG pool) were also included, purchased from JPT Peptide Technologies (Berlin, Germany).

### Assessment of ORF1p immunogenicity

Donor-matched monocytes and T cells from six healthy donors were purchased from Stemcell Technologies (Cambridge, MA). Monocytes were differentiated into dendritic cells (DCs, details in extended methods). After seven days, DCs were counted and evenly divided into ten groups per donor. Peptide pools as described above were incubated with DCs for 2 hours. T cells were thawed, then rested overnight, after which viable cells were counted, divided, and combined with DC cultures at a ratio of 10 T cells per DC, except in donor B, due to insufficient cell recovery. Cells were co-cultured for 21 days in G-rex 24-well plates (Wilson Wolf, New Brighton, MN) at 37°C and 5% CO_2_ with RPMI media (Life Technologies, Co., Carlsbad, CA) containing 10% human serum (Sigma-Aldrich, Inc., St. Louis, MO), 1M HEPES, 1x Glutamax, 20 IU/mL IL-2, 20 ng/mL IL-7, and 20 ng/mL IL-15 (all cytokines from Stemcell Tech). Every 2-3 days, 50% of media was exchanged and cytokine supplements re-added at the final concentrations above. On Day 7 of co-culture, T cells were restimulated with DCs that had been similarly derived from monocytes and pulsed with peptide pools. Supernatant samples were taken on Days 7, 11, 14, 18, and 21 and stored at –20°C until analysis. Cell samples were taken on Days 11, 14, 18, and 21 and immediately prepared for flow cytometry analysis.

Values in treatment conditions were normalized to the corresponding negative control. For significance tests, we used two-sided paired *t*-tests between values in the treatment conditions and the matched values in the negative control.

### Flow Cytometry

Briefly, cells were washed and stained with Live/Dead Viability Dye (Biolegend, San Diego, CA) and incubated on ice, in the dark, for 20 minutes. After washing, cells were blocked with human Fc block (Biolegend) prior to staining with a cell surface marker antibody cocktail specific for CD44, CD11b, CD3, 4-1BB, PD-1, CD4, CD8, CTLA-4, LAMP-1, CD154, CD69, LAG-3, CD11c, and CD25 (Biolegend). All antibodies were used between 1:100 and 1:200 dilution and incubated on ice, in the dark, for 30 minutes. After washing, cells were fixed and permeabilized using the Transcription Factor Fixation/Permeabilization kit (eBioscience, Waltham, MA) per the manufacturer’s protocol. Here, anti-FoxP3 was added at a concentration of 1:100 in 1x perm buffer for 30 minutes, on ice, in the dark. Cells were washed, resuspended, and acquired on the Cytek Northern Lights flow cytometer (Cytek Biosciences, Fremont, CA). Results were analyzed using FlowJo (Becton Dickinson, Ashland, OR) analysis software. Gating was performed utilizing fluorescence-minus-one controls.

## Results

### Analysis of large-scale public datasets reveals tumor-specific LINE-1RNA expression

Multiple studies that analyzed data from The Cancer Genome Atlas (TCGA) and similar databases have shown that LINE-1 is overexpressed and retrotranspositionally active across a variety of tumor types, with highest expression and activity in esophageal cancer (13,30).

To validate LINE-1 tumor-specificity at the RNA level, we quantified its expression across TCGA, the Cancer Cell Line Encyclopedia (CCLE), and the Genotype-Tissue Expression (GTEx) projects using L1EM, a machine learning algorithm developed expressly to address the challenges in quantifying repeat element expression in RNA-seq data (22). Locus-level accuracy in quantification is particularly important, as there are approximately half a million LINE-1 copies scattered throughout the human genome, but the vast majority are incapable of translating protein. Thus, limiting RNA quantification to only actively translating loci is paramount to determining which tumors express the target protein of interest. We took as the set of active LINE-1 loci the “intact” LINE-1 loci defined in L1Base2 (23). These are N=146 loci curated to be full length and have both intact ORF1p and ORF2p proteins in humans.

Applying L1EM to TCGA, CCLE, and GTEx data and summing across the intact LINE-1 loci revealed low expression in normal tissues (median=2.6 TPM), but high LINE-1 expression in many tumors (>25 TPM in 16% of tumor and cell line samples), including most esophageal cancer samples and consistent with previously published results (13,30) (Figures 1A, S1A). The normal (non-tumor) tissue with the highest expression of LINE-1 is testis, with an average expression of 8.1 TPM (stdev = 2.2) (Figures 1A, S1C). While ORF1 expression has been observed in the testes (31), testis is an immune-privileged tissue (32) where poor RNA/protein correlations have been observed across a variety of genes (33), so it may not yield an accurate depiction of normal tissue expression. The second highest expressing normal tissue identified in the GTEx data, esophagus – mucosa is likely a more relevant normal tissue for tumor/normal comparisons, with an average expression of 6.9 TPM (stdev = 3.3) (Figures 1A, S1C). The highest tumor expression of LINE-1 was in esophageal cancer, with 80% of samples having expression > 20 TPM (mean = 57.0 TPM, stdev = 44.9). This includes both esophageal adenocarcinoma (ESAD) (mean = 42.9 TPM, stdev = 37.8) and esophageal squamous cell carcinoma (ESCC) (mean = 74.1 TPM, stdev = 42.9).

**Figure 1.**
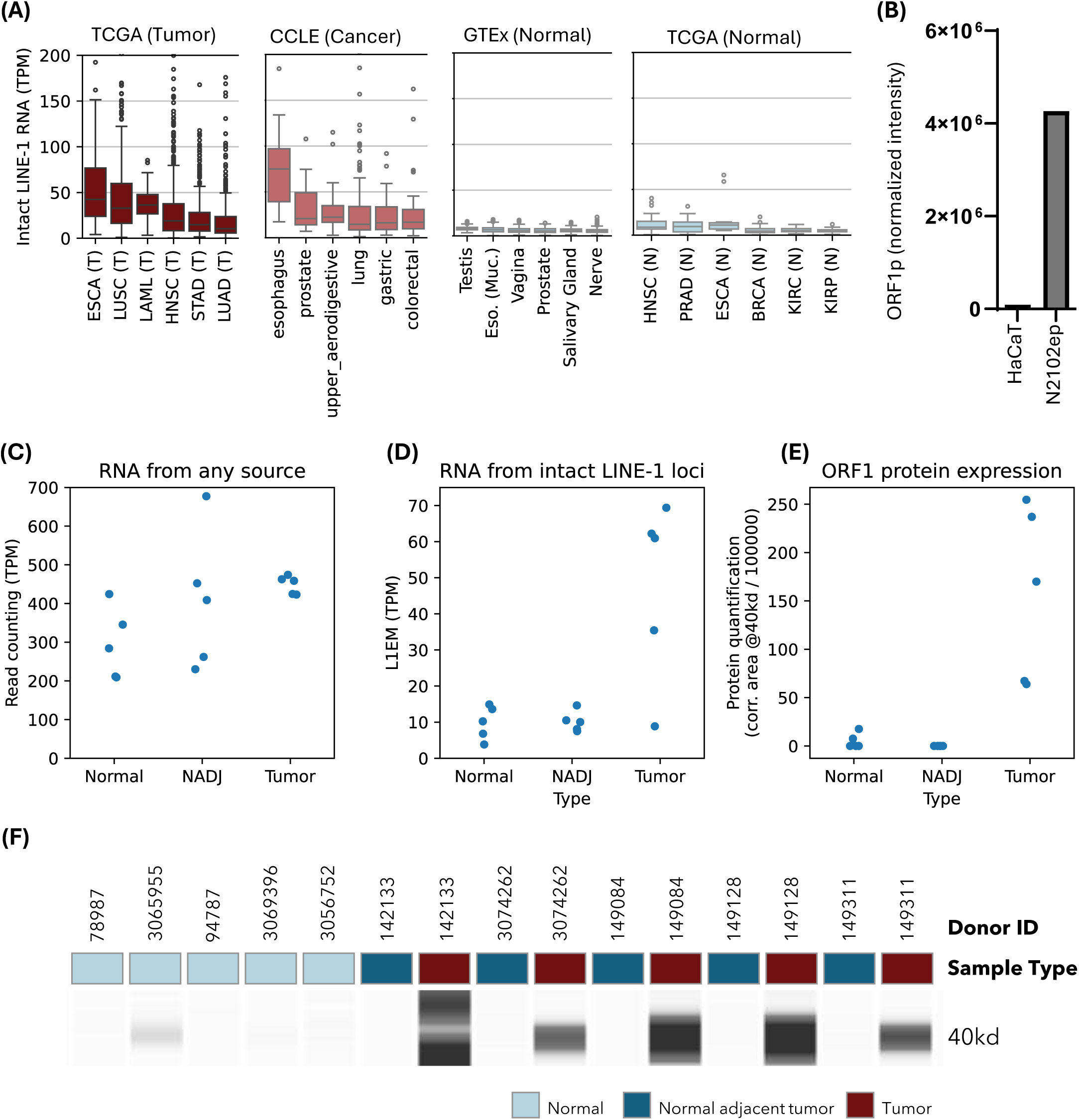
Expression of LINE-1 ORF1 in tumor and normal. (A) Active intact LINE-1 expression in top 6 TCGA tumor types, top 6 CCLE cell line tissues of origin, top 6 GTEx normal tissues and top 6 TCGA normal tissues respectively. TCGA abbreviations: https://gdc.cancer.gov/resources-tcga-users/tcga-code-tables/tcga-study-abbreviations. (B) Simple western quantification of ORF1p in HaCaT and N2102ep cells. (C) Naïve read counting quantification of L1Hs elements in esophageal cancer samples. Normal = Esophagus from individual without cancer; NADJ = Normal esophagus from a patient with esophageal cancer; Tumor = Esophageal tumor sample. (D) As in C, but quantification specifically for active expression of intact LINE-1 transcripts. (E) Quantification of ORF1 by Simple Western. (F) Image of the Western blot data summarized in panel D.

Compared to expression levels in certain tumors, the level of RNA expression of LINE-1 across all normal tissues was low—however, it was notably not zero. Since esophageal cancer is the tumor type with highest LINE-1 expression and esophagus is the non-privileged tissue with highest normal expression, we opted to assess tumor selectivity of ORF1p protein expression in esophageal cancer tumor/normal pairs.

### ORF1p protein expression is highly tumor specific in a set of esophageal tumor, tumor adjacent normal, and healthy esophagus tissue

Previous studies using immunohistochemistry (IHC) to compare LINE-1 ORF1p expression between tumor and normal samples have found its expression to be tumor specific (16,34). However, given the non-zero normal LINE-1 RNA expression observed in our transcriptomics analysis, we sought to validate the tumor specificity of ORF1p at the protein level.

We first assessed ORF1p expression in two cell lines using the Jess Simple Western (see Methods) system: N2102ep, a cell line known to express a high level of endogenous LINE-1 (35), and HaCaT, a normal-like cell line derived from human keratinocytes. Applying L1EM to RNA-seq data from these cell lines, we found that N2102ep cells have high RNA expression of LINE-1 (169 TPM) while HaCaT cells show low LINE-1 RNA expression (3.8 TPM). Consistent with our expectations and with the RNA-seq quantifications, we found high ORF1p expression in the N2102ep cells, but little to no expression detected in the HaCaT cells (Figure 1B).

The tumor specificity of ORF1p in human tissue samples was then validated. We acquired 15 tissue samples: 5 esophageal squamous cell tumor samples (ESCC), 5 matched tumor adjacent normal esophageal samples, and 5 normal esophageal samples from individuals without cancer (see Supplementary Table 1). We performed Jess Simple Western on all 15 samples. This revealed clearly discernable ORF1p expression bands in all 5 tumor samples. A faint signal was visible in one normal (see donor ID 3065955 in Supplemental Table 1) but was otherwise absent from the other 9 normal esophagus samples (Figures 1E,F), showing a highly tumor specific ORF1p expression pattern (mean 63x higher chemiluminescence in tumor than normal, p = 8x10^-5^ two-sided t-test.

### Repeat-aware quantification approaches for RNA-seq data are required to accurately predict repeat protein expression

Given the notable challenges around quantifying actively translating LINE-1 from RNA-seq data, and the fact that large, high-quality oncology omics datasets are still primarily limited to RNA-seq, we sought to validate the importance of using a locus specific approach to specifically quantify loci with intact open reading frames. To do this, we further quantified LINE-1 RNA expression in the 15 esophageal cancer and normal tissue samples and a subset of CCLE cell lines for which proteomics data exists, using a “repeat-naïve” approach, and compared these with the L1EM and Western blot protein quantification results. The naïve approach simply counts L1Hs (the evolutionarily youngest subset of L1 loci) aligned reads, without considering whether they come from transcripts likely to be translated.

In the ESCC samples, the repeat-naïve approach to quantifying LINE-1 resulted in poor differentiation of LINE-1 RNA expression levels between tumor and normal (Figure 1C) compared to L1EM (Figure 1D). Without a locus/repeat aware method like L1EM, RNA derived from the vastly greater number of inactive loci swamps any signal from protein coding LINE-1 expression. This effect has previously been described using *in silico* simulations (13,22). However, the protein quantification showed even greater tumor specificity (Figure 1E), suggesting either that not all intact loci are translated or that there may still be some residual inactive LINE-1 expression being captured by L1EM.

The need for quantification that specifically targets intact loci is further reflected in strong RNA/protein correlations from L1EM that are weaker or absent when using the naïve approach (Figure 2). For the naïve approach Pearson RNA/protein correlations are r = 0.31 (p = 0.26) for the ESCC samples (Figure 2A) and r = 0.53 (p = 0.001) for CCLE (Figure 2C). When using L1EM, the correlations are r = 0.85 (p = 5.6x10^-5^) for the ESCC sample (Figure 2B) and r = 0.8 (p = 9.3x10^-9^) in CCLE Figure 2D).

**Figure 2.**
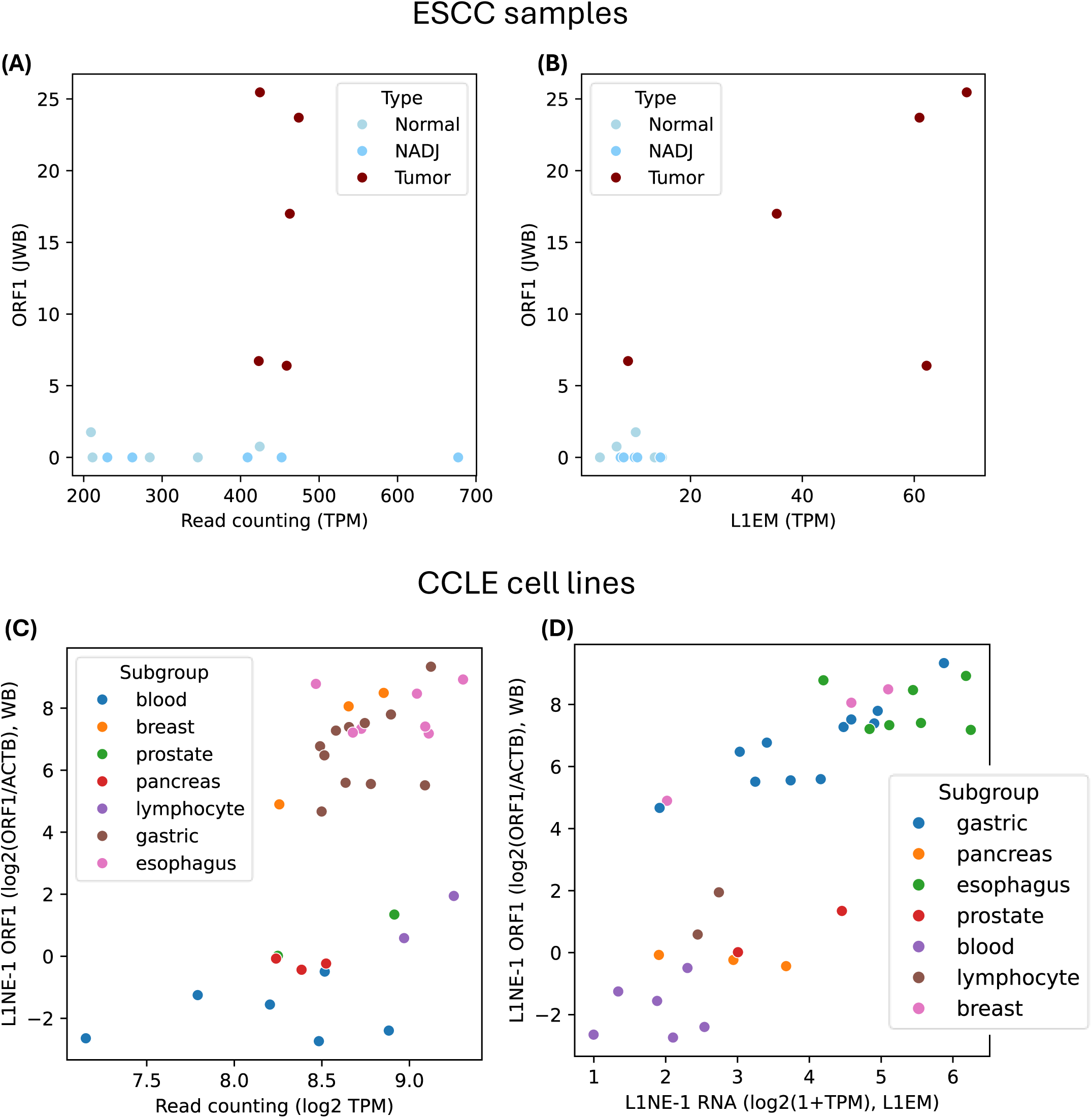
LINE-1 RNA vs ORF1 protein correlations. (A) Naïve quantification of LINE-1 (counting all L1Hs aligned reads) in esophageal tumor and normal samples compared to ORF1 protein quantification by Simple Western. (B) As in A, but using L1EM to quantify intact LINE-1 RNA expression. (C) Naïve quantification of LINE-1 in select CCLE cell lines compared ORF1 protein quantification by standard Western blot. (D) As in C, but using L1EM to quantify intact LINE-1 RNA expression.

### ORF1p peptides are detected in a comprehensive set of public immunopeptidomics data

Expression, even at the protein level, is not sufficient for identifying promising targets for a vaccine, as peptides derived from proteins must also be presented on the cell surface by MHC molecules. Further, significant population-level variability exists in terms of peptide presentation ability depending on an individual’s human leukocyte antigen (HLA) alleles, with allelic distributions varying by geography and ethnicity, among other contributing factors. Therefore, assessing peptide presentation is essential.

To understand the range of epitopes that are presented from ORF1p across tumor types and populations, we identified 47 studies with publicly available immunopeptidomics data. These data include over 1000 tumor and normal samples (Figure 3A) and cover 13 tumor types (Figure 3B). See Supplemental Table 3 for a full list of the data. We re-analyzed these data to search for ORF1p epitopes and identified 60 distinct 8, 9, 10 or 11mer epitopes matching the ORF1p consensus sequence (uniprot L1RE1). See Supplementary Table 4 for the full list of ORF1p peptides detected. ORF1p peptides spanned most of the tumor types analyzed and were largely tumor specific, with at least one ORF1p peptide detected in 19% of tumor samples, but only 4% of normal samples (Figure 3C).

**Figure 3.**
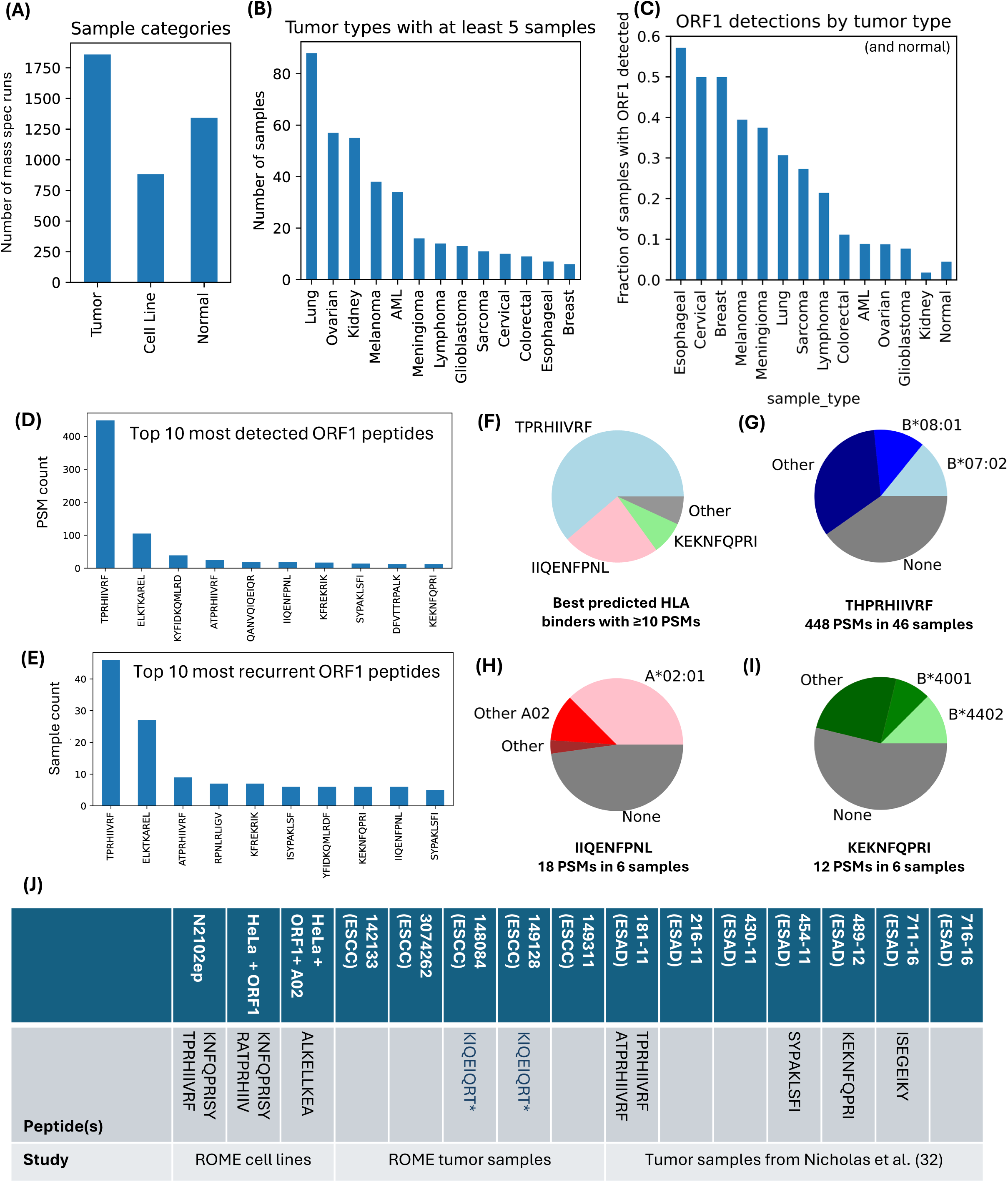
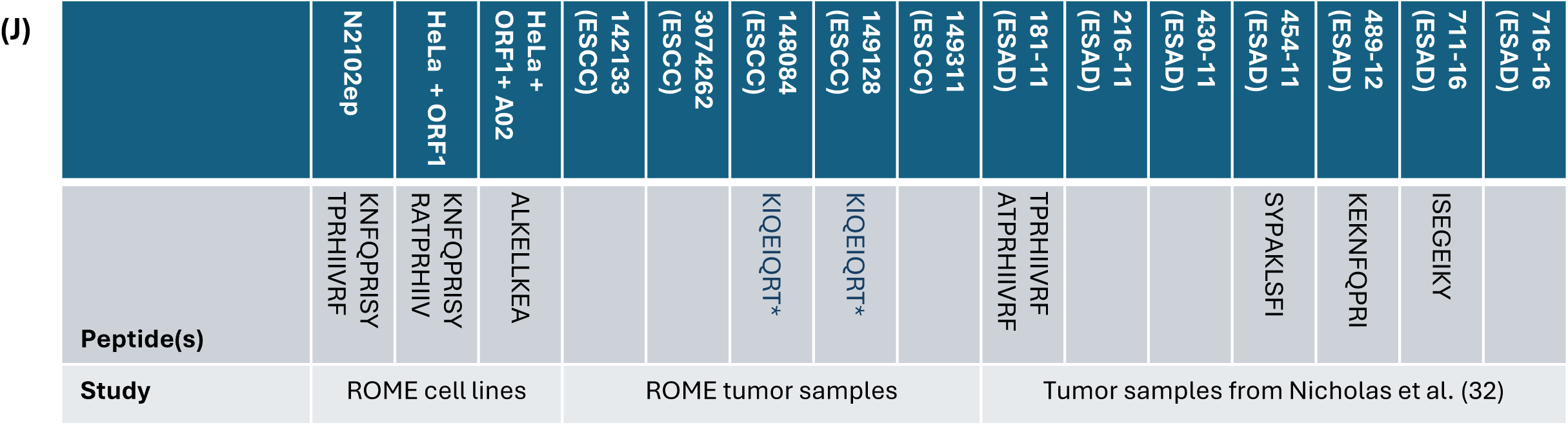
Identification ORF1p peptides in immunopeptidomics data. (A) Number of tumor, cell line and normal sample mass spectrometry runs considered. (B) Number of samples from tumor types for which at least 5 samples were considered. (C) For the tumor types shown in B, the fraction of samples where an ORF1p peptide was detected. The corresponding value for normal samples is shown for comparison. (D) The 10 most frequent peptides by number of peptide/spectrum matches (PSMs). (E) The 10 most frequent peptides by number of samples where the peptide was detected. (F) Fraction of samples predicted to present one of three frequent peptides predicted to be presented on non-overlapping alleles. Samples predicted to present more than one of these peptides are colored according to the more abundant peptide. (e.g. samples predicted to present both TPRHIIVRF and IIQENFPNL fall within the TPRHIIVRF wedge.) (G) Tumors predicted to present TPRHIIVRF broken down by the HLA allele that is expected to present that peptide. (H) As in G, but for IIQENFPNL. (I) As in G but for KEKNQPRI. (J) ORF1p peptides detected in immunopeptidomics data generated as part of this study along with detections in a publicly available study of esophageal cancer (Nicholas et al.)

Several of the detected peptides were highly recurrent, making them of particular interest (Figures 3D,E). By far the most abundantly detected peptide was TPRHIIVRF, which was detected 448 times across 46 unique samples. In particular, it is highly detected in samples with the HLA B*07:02 allele; across all tumor samples where TPRHIIVRF was detected and the HLA-B alleles are known, 88% of samples were HLA B*07:02 positive.

### LINE-1 ORF1p peptides are presented by a range of HLA alleles, making ORF1p an attractive target for a population-based vaccine

While an antigen that is highly expressed and presented by at least a narrow fraction of HLA alleles could be a development candidate, the most attractive therapeutic vaccine targets are larger proteins that are processed into multiple peptides presented on HLA alleles spanning a diverse population.

We assessed the potential of LINE-1 ORF1p to be a population-based vaccine antigen by evaluating the potential HLA alleles that would be predicted to present any of the 60 ORF1p-derived peptides that we detected in the public immunopeptidomics data, using MHCflurry (29) (a deep learning algorithm that scores the potential that a given peptide will be presented on a given allele) to score the potential of ORF1p peptides to bind the alleles reported to be present among esophageal cancer donors in the TCGA database (36).

Considering peptides that scored in the top 0.5% for a given allele (a “strong binder”), each esophageal tumor represented in the TCGA database is estimated to present at least 4, and a median of 9, ORF1p peptides. 90% of the esophageal tumors in the TCGA database harbor HLA alleles that are predicted to present at least one of three repeatedly detected peptides (THPRIIVRF, IIQENFNPL and KEKNFQPRI) (Figure 3F). TPRHIIVRF is predicted to be presented on a range of HLA-B alleles, including the two most prevalent B alleles: 07:02 and 08:01 (Figure 3G). IIQENFPNL is predicted to be presented on A*02:01, the most prevalent allele in the Western Caucasian population (Figure 3H), and KEKNFQPRI is predicted to be presented by a set of B alleles (including B*44:01 and B*40:01) that are not predicted to present THPRIIVRF (Figure 3I).

### Immunopeptidomics experiments confirm canonical and non-canonical ORF1p peptides in cell lines and tumor tissue

While re-analysis of publicly available data is a powerful and efficient way to identify ORF1p peptides, those data are not geared towards tumor types with high LINE-1 expression. It is also difficult to compare expression among tumor types, since the detection of a peptide in the sample depends both on the abundance of the peptide and the sensitivity of a given mass spectrometry experiment. To address these issues, we performed our own immunopeptidomics experiments to validate ORF1p peptide presentation in relevant cell lines and tumor tissues known to express a high level of ORF1p.

Starting with cell lines expressing high LINE-1, we generated immunopeptidomics mass spectrometry data from N2102ep and HeLa cells overexpressing LINE-1 through the recoded ORFeus construct (37). We were able to detect consensus ORF1p peptides in both cases (Figure 3J). The high prevalence of HLA-A02:01 in the Western Caucasian population makes peptides presented on that allele of particular interest. We therefore performed additional immunopeptidomics on HeLa cells overexpressing both ORF1p and HLA-A02:01. This yielded the peptide ALKELLKEA (Figure 3J), which is predicted to be a strong A02:01 binder (top 0.14% according to MHCflurry, less than the 0.5% cutoff described above), but is not among the peptides we identified in the public data.

We also generated immunopeptidomics data from the 5 esophageal cancer tissue samples described previously and searched for ORF1p consensus peptide sequences. Given that tumor samples are a mixture of cell types, with only the malignant cells expected to present ORF1p, peptides eluted from these samples were split into 5 fractions to improve sensitivity. Despite this, we did not detect consensus ORF1p peptides in these samples. This was surprising given robust detection of LINE-1 RNA and ORF1p in these samples. We therefore supplemented the search with the specific ORF1p sequences derived from the specific loci that are expressed in these samples, as predicted by L1EM. This analysis yielded an ORF1p peptide (KIQEIQRT) that was found in 2 of the 5 samples (Figure 3J). KIQEIQRT was not predicted to be strong HLA ligand by MHCflurry, but quality of the spectral match was nevertheless high (spectral angle between observed and the Spectronaut prediction was 0.88, figure S2B). The challenges in detecting ORF1p peptides contrasted with publicly available esophageal cancer data analyzed as described in the section above (figures 3C and J.)

### LINE-1 ORF1p peptides are immunogenic, eliciting interferon gamma production across multiple donors *in vitro*

To determine if ORF1p peptides are immunogenic, we used pools of 15mer peptides overlapping by 11 bp (38) to perform an *in vitro* vaccination assay using T cells and monocyte-derived dendritic cells from 6 healthy donors. Overlapping peptides spanning MOG and common viral and bacterial antigens were used as negative and positive controls, respectively. We monitored cell proliferation and interferon gamma (IFN-γ) production over time and normalized the data relative to the negative control. Cellular proliferation was detected in all conditions, with no discernible differences between ORF1p-stimulated cultures and controls (Figures 4A (log scale), S3A (absolute scale). We therefore focused on IFN-γ as a measure of a T cell-mediated immune response.

**Figure 4.**
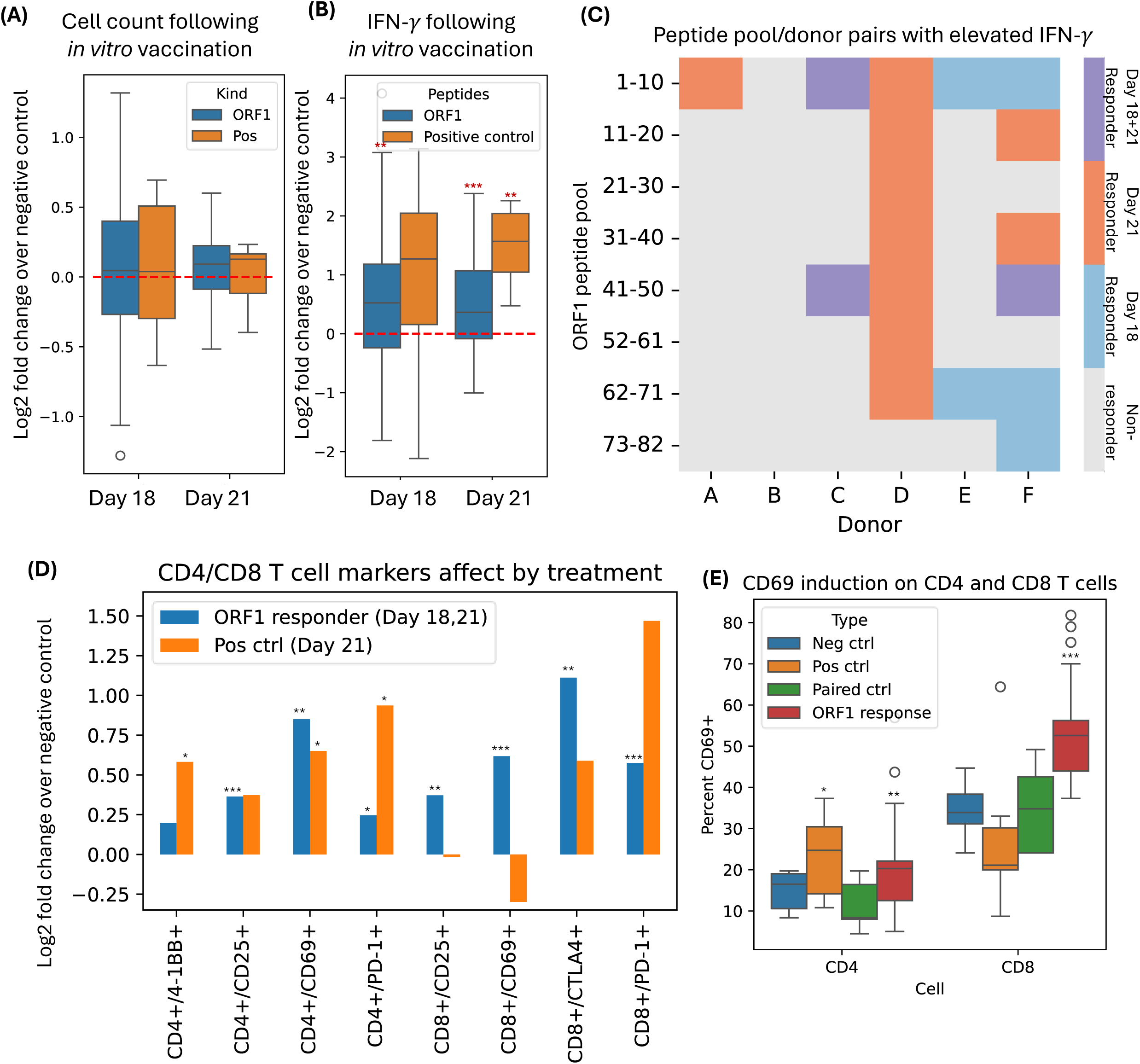
Immunogenicity of ORF1p peptides. Across all panels, paired t-test p values comparing treatment to negative control are indicated as follows: ***:p<0.001, **:p<0.01, *:p<0.05. (A) Cell counts normalized to negative control at the latter time points (day 18 and day 21) for ORF1p and positive control peptides. (B) As in A, but showing interferon gamma normalized to negative control. (C) Donor/peptide pool pairs that show interferon induction at day 18 or 21 (referred to as responders). (D) Flow cytometry markers that are significantly (p<0.05) elevated in ORF1p responders or positive control. (E) Fraction of CD4 and CD8 T cells that express the activation marker CD69 across the negative control, the positive control, the negative controls matched to the responding pairs (highlighted in panel B) and the responding pairs themselves.

Overall, peak responses varied by donor and time point. For the positive control, as shown in Figure 4B, IFN-γ production was elevated on Days 18 and 21, reaching statical significance on Day 21 (paired t-test p-value = 0.0036). While the sample size coupled with different kinetics of response per donor, did not provide sufficient power for statistical comparison of individual ORF1p peptide pools to the negative control after multiple test correction, the combined analysis of response to ORF1p stimulation was highly significant compared with the negative control (Day 18 paired t-test: 0.0064; Day 21 paired t-test: 0.00048), but was smaller than the response to the positive control (Day 18 paired t-test: 0.05; Day 21 paired t-test: 1.7x10^-8^) (see Figures 4B, S3B).

Next, to evaluate whether immune responses could be attributed to different regions of the ORF1 protein, particular donor-pool pairs were compared. Given that we expected values for non-responding donor-pool pairs to be centered around 0, we took the absolute value of the most negative value (1.8 for Day 18 and 1 for Day 21) as a cutoff to define putative responders and non-responders. These pairs are shown in Figure 4C. Peptide pool 1-10 elicited the most frequent response across donors (5 of 6 donors responded in at least one time point). Only donor B did not reach the response cutoff with any pool but trended toward increased IFN-γ production in response to pool 1-10 at Day 18 (Figure S1B). Three of six donors responded to peptide pool 62-71 for at least one time point.

### *In vitro* immune response to ORF1 includes both CD4+ and CD8+ T cells

We then assessed the specific types of immune cells that were activated during the *in vitro* vaccination assay using flow cytometry. We looked for elevated cell surface markers on CD4^+^ or CD8^+^ T cells on Day 21 for the positive control or in the ORF1 responders (mix of Day 18 and Day 21) (Figure 4D shows markers significant at p<0.05 for at least one comparison). For the ORF1p responders, we found three markers (CD25, CD69 and PD-1) that were expressed on a greater fraction of both CD4^+^ and CD8^+^ cells. This suggests that ORF1 peptides can elicit a response from both CD4^+^ and CD8^+^ T cells. In the positive control (CEFT), a greater proportion of CD4^+^ T cells expressed 4-1BB, CD69 and PD-1 compared to the negative control.

Of particular interest is the increased proportion of CD69+ T cells, a classical early marker of T cell activation (39). For the positive control, we found an increased prevalence of CD69 on CD4^+^, but not CD8^+^ T cells (p = 0.02 and 0.35, respectively, paired t-test; Figure 4D). However, for the responding ORF1p donor-pool pairs, we found increased CD69 expression on both CD4^+^ or CD8^+^ T cells (p = 0.0005 and 4x10^-6^ respectively, paired t-test; see Figure 4E). CD25 and PD-1, which can also be markers or T cell activation, were induced on T cells upon ORF1p antigen stimulation as well.

### ORF1 immunogenicity is not predicted by proposed peptide binding affinity

Given that differences in HLA alleles can lead to significant differences in immune response, we investigated whether the immunogenicity results could be predicted by the HLA type of the donors. We performed HLA genotyping of the six donors and used MHCflurry to estimate donor-pool pair affinity as follows: For each peptide pool, we took all possible 8, 9, 10, and 11mers that could be derived from the 15mers in the pool and scored their peptide-HLA affinity for each donor. The score of the best *k*-mer in each donor-pool pair is shown in Figure 5A. Notably, while there is significant variation in the scores, even the donor-pool pair with lowest affinity score (pool 1-10 and donor C) still includes an ORF1 peptide (TEQSWMENDF) that scores in the top 0.75% for one of Donor C’s alleles (B*44:02). Given the prediction of near universal presentation across donor/pool pairs, it is perhaps not surprising that we did not see a correlation between predicted presentation and interferon production (Figure 4B, r = 0.11, p=0.45).

**Figure 5.**
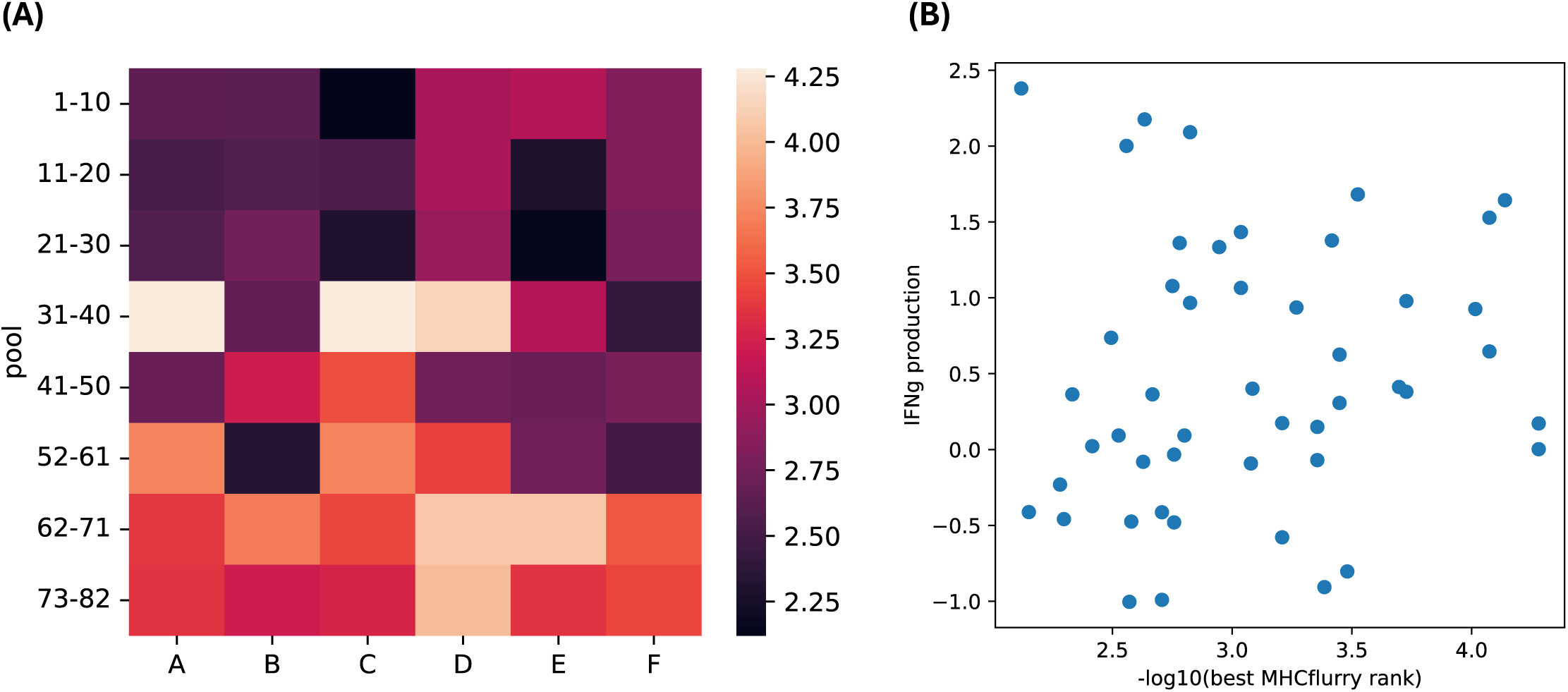
MHCflurry presentation prediction for in vitro vaccination donors. (A) -Log10 of the rank of the best peptide presentation prediction among 8-11mers in that pool on alleles present in that donor. (B) Comparison of values shown in panel A to interferon gamma induction.

## Discussion

Over the last several years, there has been an explosion of new cancer vaccine modalities and delivery systems. However, to be effective, a good cancer vaccine requires both an adequate delivery system and an immunogenic antigen. LINE-1 ORF1p has long been known to be highly expressed in multiple tumor types (15,16), but it has been underexplored as a tumor antigen, perhaps because LINE-1 is a highly repetitive element that is challenging to quantify at the RNA level.

Here we show how repeat-aware quantification algorithms reveal tumor specificity of LINE-1 at the RNA level and validate this at the protein level in a set of esophageal tumor and normal samples. In a comprehensive meta-analysis across 47 public immunopeptidomics data sets, spanning 13 tumor types, we identify ORF1p peptides present in tumor types beyond esophageal cancer, including breast, cervical, and lung cancers, and melanoma. We then performed our own immunopeptidomics experiment in cell lines and tumor tissues expressing ORF1p. While we did validate presentation in the cell lines, validation in tumor tissues was more challenging, likely due to lower tumor purity and the limited sensitivity of mass spectrometry proteomics. Our results may also be impacted by difference in population (our samples were of east Asian descent whereas most of the public data comes from a Caucasian population.) Finally, we used an *in vitro* vaccination assay to assess the ability of T cells from healthy donors to respond to peptides derived from ORF1p and found that selected donor-peptide pairings led to increased IFN-γ production. The T cells cultured represent only a small portion of the donor’s T cell repertoire. Therefore, the fact that we observed a response in most donors, even with a potentially stringent cutoff, is promising. This supports the idea that while LINE-1 is encoded in the genome, it is not fully recognized as self, making it a potential vaccine target.

We also used flow cytometry to dissect the T cells in the responding peptide-donor pairs and found evidence for activation of both CD4 and CD8 T cells. This is key as both types of T cells are critical for an optimal antitumor immune response (40). However, even when T cells are activated, this alone does not necessarily indicate cytotoxicity. Future work on ORF1p as a cancer antigen will be needed to assess the cell killing capabilities of T cells targeting ORF1p, *in vitro* and/or *in vivo*.

This work highlights the potential of LINE-1 ORF1p as a cancer vaccine antigen. It is widely and specifically expressed across tumors. Peptides derived from ORF1 are frequently presented on MHC class I molecules in tumors, span a range of HLA alleles, and can activate both CD4^+^ and CD8^+^ T cells. In this study we highlighted expression in esophageal tumors, but numerous studies have highlighted a relationship between p53 mutation and LINE-1 activation (13,30,41–44). There are also several studies showing that LINE-1 activation occurs in early tumorigenesis (16,45,46). Therefore, an efficacious ORF1p vaccine could be relevant to a range of solid tumors and perhaps even for onco-prevention.

## Declarations

### Ethics approval and consent to participate

This study used publicly available data, established cell lines and retrospectively collected samples only. Esophageal cancer patients were consented for sample collection and analysis. Consent forms were provided to ROME Therapeutics by BioIVT upon purchase of samples.

### Consent for publication

All authors have consented to publication.

### Availability of data and materials

Mass spectrometry data considered in the study is available on the proteomexchange (https://www.proteomexchange.org); see table S3 for a full list of accession numbers. TCGA and GTEx data were accessed (and are available) through the Terra platform (https://terra.bio).

### Author’s contributions

WM performed computational analyses, oversaw work performed at CROs, contributed to study design, and worked on the manuscript draft. MS performed and designed in vitro vaccination assay. SC contributed to design of computational analyses and performed additional computation analyses. DK performed additional lab work. HK provided expertise, set direction and contributed to study design. LD provided project oversight and expertise, contributed to study design and led manuscript preparation.

## Supporting information

Extended Methodss

## Acknowledgments

The authors acknowledge lab support and oversight from Jared Steranka, Nicole Van Hoeken and Kimberly Long. The study authors benefited greatly from immunological and cancer vaccine expertise provided by Jessica Baker Flechtner. This project grew out of work designed by the authors along with Menachem Fromer and Dennis Zaller and performed by the authors along with Bryan Thornlow.

## List of abbreviations

TAA: Tumor-Associated Antigen
MHC: Major Histocompatibility Complex
TCGA: The Cancer Genome Atlas
GTEx: Genotype-Tissue Expression
CCLE: Cancer Cell Line Encyclopedia
TPM: Transcripts Per Million
DDA: Data Dependent Acquisition
PSM: Peptide Spectral Matches
FDR: False Discovery Rate
Da: Dalton
EBV: Epstein-Barr Virus
HCMV: Human Cytomegalovirus (HCMV)
CEFT: Clostridium tetani, Epstein-Barr Virus, Human Cytomegalovirus, Influenza A
MOG: Myelin Oligodendrocyte Glycoprotein
CD: Dendritic Cells
ESAD: ESophageal ADenocarcinoma
ESCC: Esophageal Squamous Cell Carcinoma (ESCC)
IHC: ImmunoHistoChemistry (IHC)
HLA: Human Leukocyte Antigen
IFN: InterFeroN

## Supplementary Table Legends

Table S1. Esophageal sample info. Demographic, HLA and purity information for the esophageal samples analyzed in this study.

Table S2. Rome immunopeptidomes. Raw proteomics files generated as part of this study.

Table S3. Public raw files. URL, HLA, sample type and sample name information for the publicly available raw files analyzed as part of this study.

Table S4. ORF1 peptide / spectrum matches identified from the raw files listed in table S3 and FDR < 1%.

Table S5. Cell counts normalized to negative control from *in vitro* vaccination assay.

Table S6. Interferon gamma data from *in vitro* vaccination assay. Values in Day7, Day11, Day14, Day18, Day21 are raw estimates of interferon gamma in pg/mL. The following columns are normalized to the corresponding negative control.

Table S7. Flow cytometry data. (A) Legend to match rows in the following table to correct peptide pool / donor pairing. (B) Flow data collected at day 11. (C) Flow data collected at day 14. (D) Flow data collected at day 18. (E) Flow data collected at day 21.

Table S8. HLA information for the 6 donors using in the *in vitro* vaccination assay. HLA typing was done by CD genomics utilizing amplicon sequencing.

---

**Figure S1.**
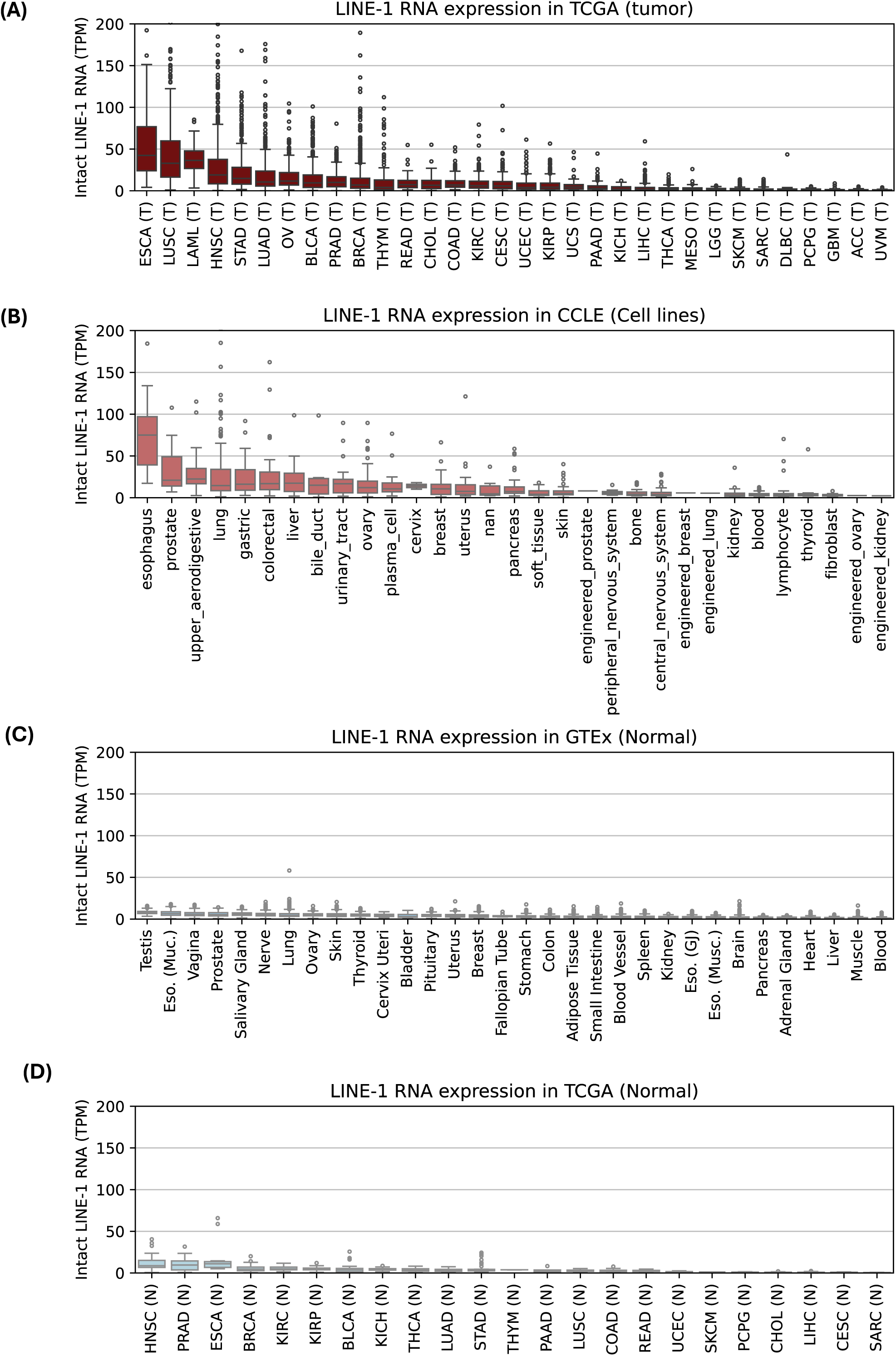
Expression of LINE-1 ORF1 in tumor and normal (extended.) (A) Active intact LINE-1 expression for all TCGA tumor types. TCGA abbreviations: https://gdc.cancer.gov/resources-tcga-users/tcga-code-tables/tcga-study-abbreviations. (B) Active intact LINE-1 expression for all CCLE cell lines. (C) Active intact LINE-1 expression for GTEx normal tissues. (D) Active intact LINE-1 expression for all TCGA normal tissues. TCGA abbreviations: https://gdc.cancer.gov/resources-tcga-users/tcga-code-tables/tcga-study-abbreviations.

**Figure S2.**
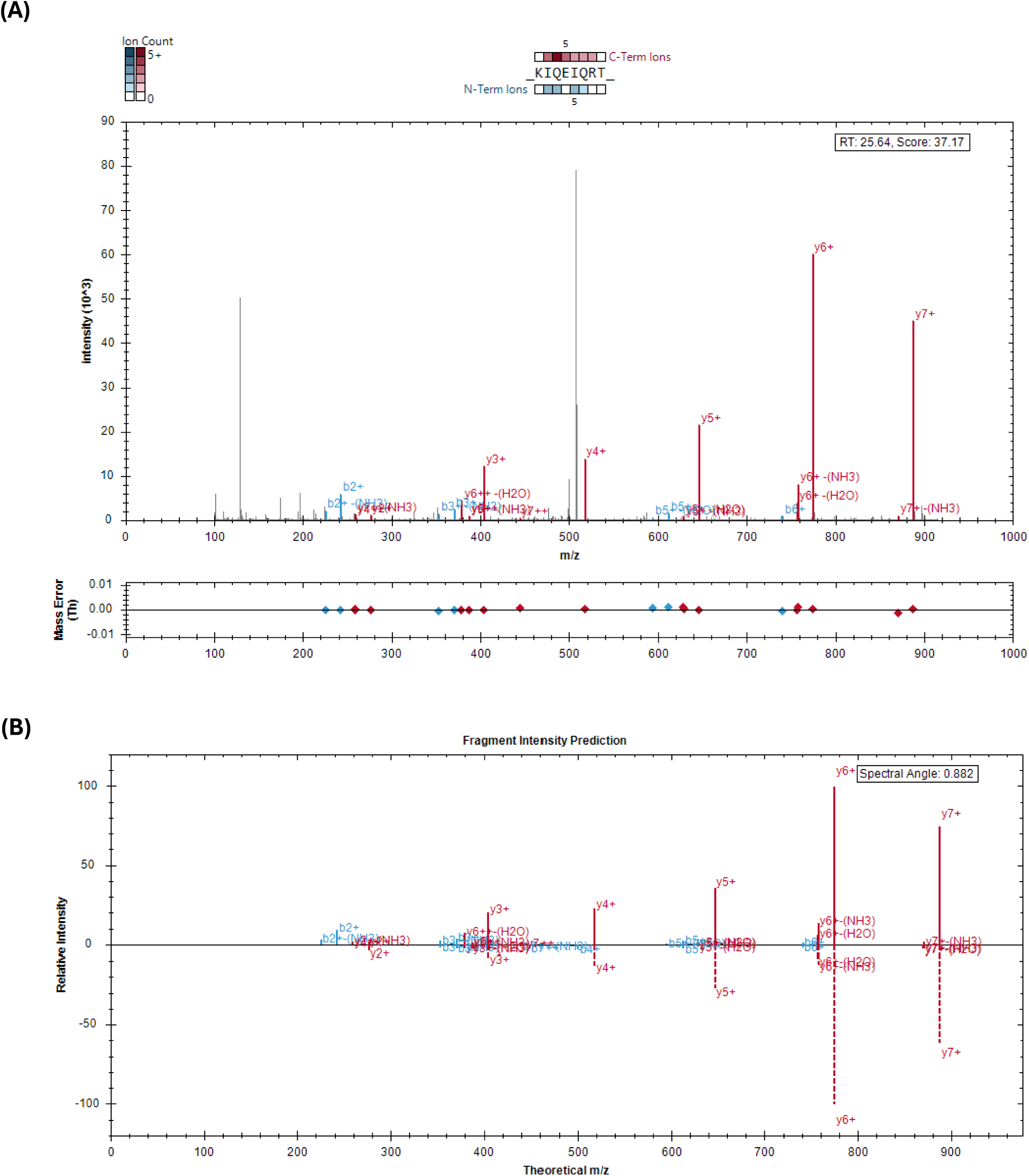
Inspection of KIQEIQRT spectrum. (A) MS2 spectrum for this peptide. Matching peptide and fragment masses are colored in green, red & blue. (B) Comparison of *in silico* predicted (dashed lines) and the experimentally observed (solid lines) fragmentation profile.

**Figure S3.**
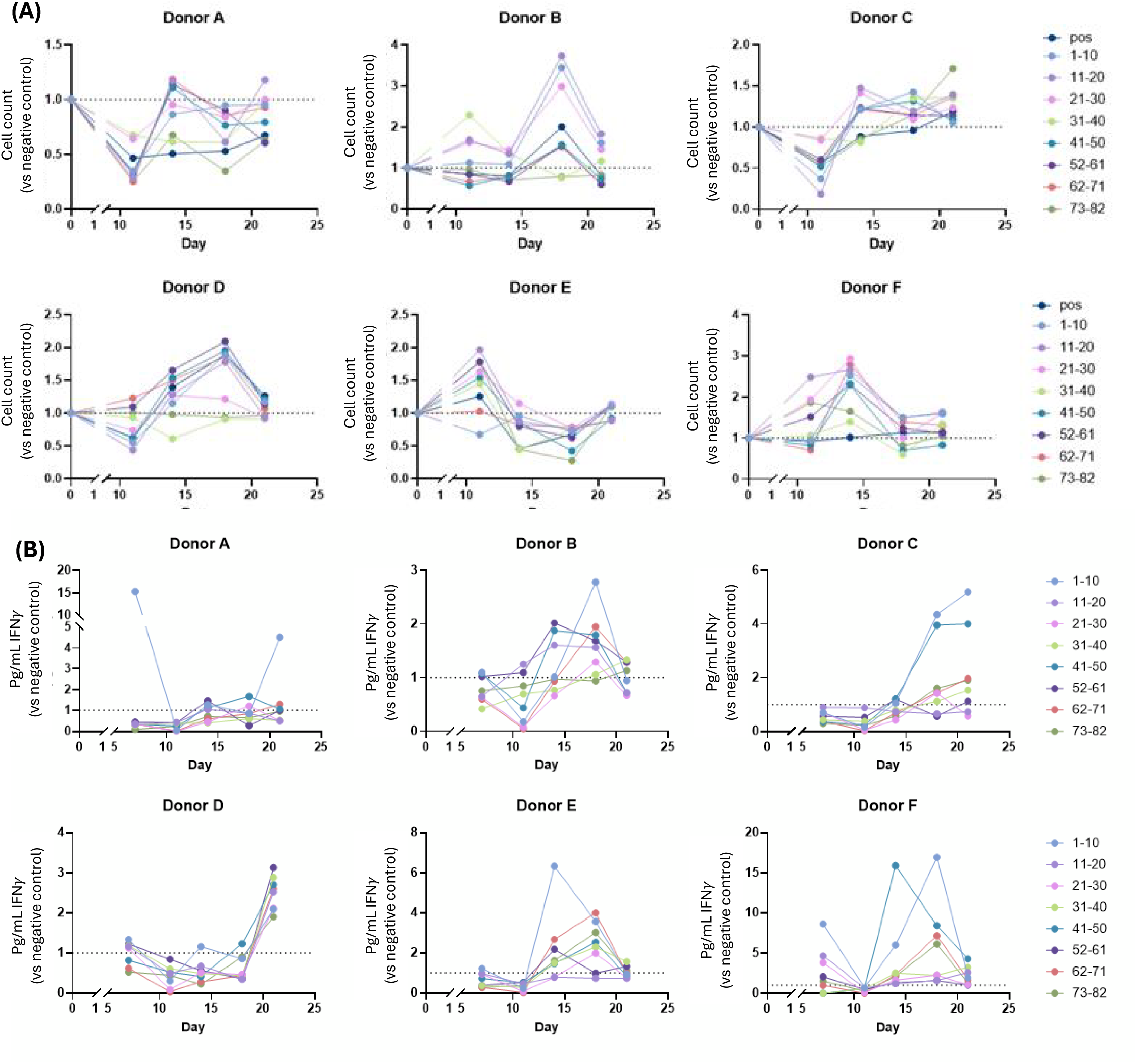
Immunogenicity of ORF1 peptides extended. (A) Time series of cell counts for each ORF1p peptide pool and each donor. Values are normalized to corresponding negative control. (B) Time series of interferon gamma production for each ORF1p peptide pool and each donor. Values are normalized to corresponding negative control.

