## Extended Methodss for "LINE-1 ORF1p is a shared and immunogenic antigen in cancer"

### Extended Methods

##### Quantification of LINE-1 from RNA-seq

Analysis was performed as in our previous study^13^. Briefly, RNA-seq reads from TCGA and GTEx were aligned to the GRCh38 human gene reference with ensemble v104 gene annotation using the STAR aligner^40^. Locus specific LINE-1 expression was quantified using L1EM^18^ from the star aligned bams. Read counts were extracted from the “full_counts.txt” output and then converted to transcripts per million (TPM) by combining with the read counts output from STAR. The TPM values were then summed across loci that overlap the “Human Full-Length, Intact LINE-1 Element [FLI-L1]” loci from L1base^19^ (<https://l1base.charite.de/BED/hsflil1_8438.bed>.) Naïve (read counting) quantification was done using samtools (https://www.htslib.org/) and the repeatmasker track on the UCSC genome browser (<https://genome.ucsc.edu/cgi-bin/hgTables?hgsid=2401253997_xg6mShcYPHVELoYFifJn3MHAqEuT>) to extract all reads with an alignment that overlaps an L1Hs element. The number of unique read ids was then counted using the “unique” command in bash and normalized to the total number of unique read ids in the bam file. Analysis and data acquisition was done using the Terra platform (terra.bio).

##### Cancer Tissue Preparation

5 esophageal tumor, 5 non-malignant adjacent, and 5 healthy esophageal samples were obtained from BIOIVT. All tumor and adjacent normal samples were from Vietnamese individuals with esophageal squamous cell carcinoma. Fully normal samples were from white residents of the United states. (Full sample details can be found in supplemental table 1). Frozen tissue was pulverized using the Covaris cryoPREP instrument (Woburn, MA).

##### Immunoblotting

For preparation of protein lysates, cryopulverized patient tumors were lysed in TPER (Thermo Fisher #78510) supplemented with protease/phosphatase inhibitor (Cell Signaling Technologies #5872). Lysates were clarified by centrifugation at 15,000 rcf for 15 minutes at 4C, and protein levels were quantified by bicinchoninic acid (BCA) assay (Pierce #A53225). Proteins were denatured in reducing Laemmli loading buffer at 100°C for 5 minutes.

##### Jess Western

ORF1p Western blots (excluding CCLE cell lines shown in figure 2 C,D) were done using the Protein Simple Jess Automated Western Blot System, using a commercially available rabbit monoclonal antibody (ab246317 from abcam). Bands from 39kd to 50kd were considered ORF1. ORF1 is a 40kd protein but sometimes runs higher in this system, especially when cell line extracts are used. The corrected area (corr. area) was used for quantification and values were normalized to 2µg of total loaded protein.

##### Traditional Western (CCLE cell lines)

Equivalent amounts of total proteins were separated on 4-15% Tris-Glycine TGx SDS-polyacrylamide gel electrophoresis (PAGE) gels (Bio-Rad #5671085). Proteins were transferred to nitrocellulose using standard methods, and membranes were blocked in blocking solution containing 5% bovine serum albumin (BSA) in tris-buffered saline (TBS) with 0.2% Tween 20. Primary antibodies against ORF1p (Abcam #ab245249), ORF1p (Sigma # MABC1152) and Cofilin (CST #5175) were diluted in blocking solution and were incubated with membranes at 4°C overnight. Horseradish peroxidase-conjugated secondary antibodies were diluted in blocking solution (1:5,000; Jackson ImmunoResearch) and incubated with membranes at room temperature for 1 hour. Western blots were developed using West Pico PLUS Super Signal ECL reagents (Thermo Fisher #34578) and imaged on an iBright CL750 (Thermo Fisher). ORF1 quantification in CCLE cells was performed at Pharmaron using a standard Western blot and a different commercially available ORF1 antibody (Sigma: MABC1152). Quantifications were normalized to actin beta.

##### HeLa overexpression systems

To generate stable HeLa cell lines expressing DOX-inducible L1 ORFeus and C-terminally tagged HLA-A*0201, piggyBac-compatible plasmids were transfected into cells along with the piggyBac transposase. The cells were then selected using Hygromycin (100 µg/ml) and Puromycin (0.5 µg/ml) for one week. Subsequently, the cells were expanded at scale, and 100 million cells were submitted for peptide immunopeptidomics studies. HeLA cells were cultured in DMEM medium supplemented with 10% Tet-free FBS and 1% Pen-Strep.

##### Immunopeptidomics experiments

Immunopeptidomics was performed by Biognosys using the TrueDiscovery® platform. Briefly, samples were lysed and MHC class I complexes were immunoprecipitated using the W6/32 antibody and peptides were eluted from the complexes. Tissue samples were split into 5 fractions; cell line samples were not fractionated. Peptides were then loaded on a liquid chromatography column and analyzed by data dependent acquisition (DDA) mass spectrometry. Peptide spectral matches (PSMs) at a false discover rate (FDR) of <1% were identified by the Spectronaut software. For the esophageal tumor samples, the search database was supplemented by ORF1s translated from loci that have an uninterrupted ORF1 and are expressed at least 2TPM (L1EM estimate) in at least one the tumor samples.

##### Re-analysis of public immunopeptidomics data

Raw files were downloaded from the relevant repository and then converted to mzml using msconvert (<https://proteowizard.sourceforge.io/download.html>). Comet was used to query mass spectra against the uniprot human proteome (<https://www.uniprot.org/proteomes/UP000005640>). Decoy search was set to concatenated search, peptide mass tolerance was set to 20ppm, fragment bin tolerance was set to 0.02 Da, search enzyme was set to none, allowed peptide length was 8-11, oxidation (15.9949 Da) of methionine was a variable modification and carboxyamidomethylation (57.021464 Da) of cysteine was a fixed modification. The comet matches were then rescored using the Prosit^41^ model as implemented in Oktoberfest^42^. Peptides that matched ORF1 (<https://www.uniprot.org/uniprotkb/Q9UN81/entry>) but no other protein were considered ORF1 peptides. ORF1 hits at an FDR of 1% were identified using a semisupervised SVM similar to the strategy outlined by percolator^43^. All of the Oktoberfest parameters were considered as features along with the best EL score from netMHCpan^44^ across a panel of common HLA alleles (listed in figure 3E). As an initial guess, PSMs with spectral_angle>0.25, count_observed_y≥5, fraction_observed_and_predicted_vs_predicted>0.5 and abs_rt_diff≤5 were select as the positive training set. Decoy PSMs were selected as the negative training set. An svm was trained on this data using sklearn python package and probability scores were calculated for the inclusion of each PSMs in the positive set. A probability score cutoff was then chosen to yield hits at a target/decoy FDR of 1%. The chosen hits were then used as the new positive training set and the process was repeated 15 times to yielding the final call set.

##### Generation of ORF1 peptides for T cell immunogenicity assay

Using the canonical peptide sequence of ORF1p, we generated a list of 82 15-mer peptides with 11 bp overlaps. Peptides were sequenced by GenScript Biotech (Piscataway, NJ). Peptides’ recovery with >90% purity were included in our immunogenicity experiment, resulting in 80 peptides moving forward to testing. Each peptide was reconstituted to 8mM DMSO and used at a final concentration of 1uM. A positive control consisting of 27 peptides from common pathogens (Clostridium tetani, Epstein-Barr Virus (EBV), Human Cytomegalovirus (HCMV), Influenza A; also known as the CEFT pool) and a negative control consisting of peptides from the human MOG gene (Myelin Oligodendrocyte Glycoprotein; also known as the MOG pool) were also included, purchased from JPT Peptide Technologies (Berlin, Germany). Positive and negative peptide pool controls also were used at final concentration of 1uM. Peptides were combined into groups of ten, in order, starting at the N’ terminus of ORF1p, resulting in eight groups for immunogenicity testing.

##### Assessment of ORF1 immunogenicity

Donor-matched monocytes and T cells were purchased from Stemcell Technologies (Cambridge, MA) from six healthy donors. Monocytes were differentiated into dendritic cells (DCs) using the Immunocult Dendritic Cell Culture Kit (Stemcell Tech), following the manufacturer’s protocol. After seven days, DCs were counted and evenly divided into ten groups per donor. Peptide pools as described above were incubated with DCs for 2h. T cells rested overnight similarly were counted, divided, and combined with DC cultures. Cells were co-cultured in G-rex 24 well plates (Wilson Wolf, New Brighton, MN) at 37C and 5% CO_2_ with RPMI media (Life Technologies, Co., Carlsbad, CA) containing 10% human serum (Sigma-Aldrich, Inc., St. Louis, MO), 1M HEPES, 1x Glutamax, 20IU/mL IL-2, 20ng/mL IL-7, and 20ng/mL IL-15 (all cytokines Stemcell Tech). Co-culture lasted 21 days. Every 2-3 days, 50% of media was exchanged and cytokine supplements re-added at the final concentrations above. On day 7 of co-culture, T cells were restimulated with DCs that had been similarly expanded from monocytes and pulsed with peptide pools. Supernatant samples were taken on days 7, 11, 14, 18, and 21 and stored at –20C until analysis. Cell samples were taken on days 11, 14, 18, and 21 and immediately prepared for flow cytometry analysis.

##### Flow Cytometry

Briefly, cells were washed and stained with Live/Dead Viability Dye (Biolegend, San Diego, CA) and incubated on ice, in the dark, for 20 minutes. After washing, cells were blocked with human Fc block (Biolegend) prior to staining with cell surface marker cocktail for CD44, CD11b, CD3, 4-1BB, PD-1, CD4, CD8, CTLA-4, LAMP-1, CD154, CD69, LAG-3, CD11c, and CD25 (all Biolegend). All antibodies were used between 1:100 and 1:200 dilution and incubated on ice, in the dark, for 30 minutes. After washing, cells were fixed and permeabilized using the Transcription Factor Fixation/Permeabilization kit (eBioscience, Waltham, MA) per the manufacturer’s protocol. Here, FoxP3 was stained at a concentration of 1:100 in 1x perm buffer for 30 minutes, on ice, in the dark. Cells were washed, resuspended, and run on the Cytek Northern Lights flow cytometer (Cytek Biosciences, Fremont, CA). Results were analyzed using FlowJo (Becton Dickinson, Ashland, OR) analysis software. All washing steps were performed twice in DPBS. Gating was performed utilizing fluorescence-minus-one controls.
